# Evaluating the interplay between estrous cyclicity and induced seizure susceptibility in *Scn2a^K1422E^* mice

**DOI:** 10.1101/2023.04.27.538584

**Authors:** Dennis M. Echevarria-Cooper, Jennifer A. Kearney

**Affiliations:** Department of Pharmacology, Feinberg School of Medicine, Northwestern University, Chicago, IL 60611, USA; Northwestern University Interdepartmental Neuroscience Program, Northwestern University, Chicago, IL 60611, USA

## Abstract

Pathogenic variants in *SCN2A* are associated with a range of neurodevelopmental disorders (NDD). *SCN2A*-related NDD show wide phenotypic heterogeneity, suggesting that modifying factors must be considered in order to properly elucidate the mechanisms of pathogenic variants. Recently, we characterized neurological phenotypes in a mouse model of the variant *SCN2A*-p.K1422E. We demonstrated that heterozygous *Scn2a*^*K1422E*^ female mice showed a distinct, reproducible distribution of flurothyl-induced seizure thresholds. Women with epilepsy often show a cyclical pattern of altered seizure susceptibility during specific phases of the menstrual cycle which can be attributed to fluctuations in hormones and corresponding changes in neurosteroid levels. Rodent models have been used extensively to examine the relationship between the estrous (menstrual) cycle, steroid hormones, and seizure susceptibility. However, the effects of the estrous cycle on seizure susceptibility have not been evaluated in the context of an epilepsy-associated genetic variant. To determine whether the estrous cycle affects susceptibility to flurothyl-induced seizures in *Scn2a*^*K1422E*^ female mice, estrous cycle monitoring was performed in mice that had undergone ovariectomy (OVX), sham surgery, or no treatment prior to seizure induction. Removing the influence of circulating sex hormones via OVX did not affect the non-unimodal distribution of flurothyl seizure thresholds observed in *Scn2a*^*K1422E*^ females. Additionally, flurothyl seizure thresholds were not associated with estrous cycle stage in mice that underwent sham surgery or were untreated. These data suggest that variation in *Scn2a*^*K1422E*^ flurothyl seizure threshold is not significantly influenced by the estrous cycle and, by extension, fluctuations in ovarian hormones. Interestingly, untreated *Scn2a*^*K1422E*^ females showed evidence of disrupted estrous cyclicity, an effect not previously described in a genetic epilepsy model. This unexpected result highlights the importance of considering sex specific effects and the estrous cycle in support of more inclusive biomedical research.

## Introduction

Genetic variants in *SCN2A*, encoding the NaV1.2 voltage-gated sodium channel, are associated with a range of neurodevelopmental disorders (NDD) including severe epilepsy syndromes and autism spectrum disorder (ASD)^1^. Phenotypic variability observed across *SCN2A*-related NDD can be partially attributed to distinct changes in the biophysical properties of mutant channels^2–4^. However, recurrent and inherited variants show heterogeneity even among individuals with the same variant, suggesting that phenotype expressivity may be subject to modifying factors^5^. The variant *SCN2A*-p.K1422E is associated with infant-onset developmental delay, infantile spasms, and features of ASD. We previously demonstrated that both male and female *Scn2a*^*K1422E*^ heterozygous mice (abbreviated as *Scn2a*^*E/+*^ going forward) had a higher threshold for flurothyl-induced GTCS compared to WT^6^. Within that initial data set, we also noted that the cumulative distribution of latencies to GTCS in *Scn2a*^*E/+*^ females was significantly different from the other groups^6^. This effect was subsequently replicated in two additional cohorts of *Scn2a*^*E/+*^ and WT females^6^. This is suggestive of a potential genotype-dependent interaction with a sex-specific modifying factor. Furthermore, sex differences have been reported in a number of NDD including epilepsy and ASD^7–9^.

Epilepsy represents a diverse group of conditions for which differences between males and females can vary across different types of seizures^8,10–12^. The neurobiological basis for sex differences in seizure susceptibility is likewise varied and the subject of ongoing research^10,11^. It has been widely documented that women with epilepsy often show a cyclical pattern of altered seizure susceptibility during specific phases of the menstrual cycle^13–15^. This pattern of “catamenial epilepsy” is a neuroendocrine condition that affects between 25-75% of women with epilepsy^16^. Additionally, menstrual disorders are more common in women with epilepsy than in the general population^17^. The association between seizure susceptibility and the menstrual cycle can be attributed to fluctuations in hormones and corresponding changes in neurosteroid levels^10,11,14,18^. Rodent models have been used extensively to examine this association, revealing complex, sometimes contradictory effects of ovarian hormones and neurosteroids on seizure susceptibility^11,19–24^. In general, estradiol is proconvulsant and progesterone (along with its neurosteroid derivative allopregnalone) is anticonvulsant^10,11,25^. Woolley and colleagues showed previously that neither estradiol nor progesterone affect susceptibility to flurothyl-induced seizures in ovariectomized rats^20^. However, it is possible that the K1422E variant may affect sensitivity to ovarian hormones. The nuclear estrogen receptor alpha is a transcription factor that is activated by endogenous estrogens (e.g. estradiol) and has been shown to target *Scn2a* in sexually dimorphic brain regions^26^. Based on these findings, we hypothesized that the estrous cycle affects susceptibility to flurothyl-induced seizures in *Scn2a*^*E/+*^ female mice.

To date, the effects of the estrous cycle on seizure susceptibility have not been evaluated in the context of an epilepsy-associated genetic variant. We applied formal analyses of modality to our previously published flurothyl data and established that the distribution of latencies to GTCS in *Scn2a*^*E/+*^ females was significantly non-unimodal. To test our hypothesis, we sought to answer two essential questions. First, we wanted to know if circulating sex hormones (i.e., the estrous cycle) are necessary to observe the non-unimodal distribution of flurothyl seizure thresholds in *Scn2a*^*E/+*^ mice. Second, we wanted to know if flurothyl seizure thresholds are associated with a particular stage of the estrous cycle. Here, we examined flurothyl seizure thresholds in *Scn2a*^*E/+*^ female mice that underwent ovariectomy or sham surgery and subsequent estrous cycle monitoring. We found that ovariectomy did not affect the non-unimodal distribution of flurothyl seizure thresholds observed in *Scn2a*^*E/+*^ mice. Additionally, flurothyl seizure thresholds were not associated with estrous cycle stage in mice that underwent sham surgery, nor in a non-surgerized cohort. These data suggest that the estrous cycle does not significantly affect susceptibility to flurothyl-induced seizures in *Scn2a*^*E/+*^ mice. However, we did find evidence of disrupted estrous cyclicity in the cohort of non-surgerized *Scn2a*^*E/+*^ mice.

## Materials and Methods

### Mice

Female heterozygous *Scn2a*^*E/+*^ and WT mice for experiments were obtained from the line *Scn2a*^*em1Kea*^ (MGI:6390565; MMRRC:069700-UCD), which is maintained as an isogenic strain on C57BL/6J (#000664, Jackson Laboratory, Bay Harbor, ME). Mice were maintained in a specific pathogen free barrier facility with a 14 h light/10 h dark cycle and access to food and water ad libitum. All animal care and experimental procedures were approved by the Northwestern University Animal Care and Use Committees in accordance with the National Institutes of Health Guide for the Care and Use of Laboratory Animals. Principles outlined in the ARRIVE (Animal Research: Reporting of *in vivo* Experiments) guideline were considered when planning experiments^27^.

### Surgeries

Bilateral ovariectomy (OVX) or sham surgery was performed on female WT and *Scn2a*^*E/+*^ mice at 5-6 weeks of age as previously described^28^. For both surgeries, subjects were deeply anesthetized with a cocktail of ketamine and xylazine (100 mg/kg and 10 mg/kg, respectively, IP), followed by administration of preoperative analgesia (20 mg/kg meloxicam, SC) and local anesthesia for the primary incision site (infiltration using 20 µL of 0.2% lidocaine). A primary midline skin incision and bilateral body wall incisions were performed to access the ovaries. For OVX, the ovarian fat pads were exteriorized, a crush injury was induced in the uterine horns below the ovaries, and the ovaries were subsequently removed. For sham surgeries, no crush injury was induced, and the intact ovarian fat pads were re-internalized following identification of the ovaries. At the conclusion of surgery, sustained release buprenorphine was administered (1 mg/kg, SC) for post-operative analgesia and atipamezole hydrochloride was administered (1 mg/kg, SC) to reverse the effects of anesthesia. All surgically treated subjects were housed individually during recovery and estrous cycle monitoring.

### Estrous cycle monitoring

Estrous cycle monitoring was performed using a vaginal lavage protocol as previously described^29^. Cycle monitoring was conducted from 0700 to 0800 daily. Vaginal cytology was assessed using light microscopy (Olympus CX33 Biological Microscope) and representative images of each sample at 10x magnification were captured using an Apple iPhone 13 Pro (Camera app; 3x zoom) via the microscope eye piece. Estrous cycle stage was determined by evaluating the proportion of relevant cell types in a given sample as previously described^30–32^. An additional classification, “unclear” was used to designate samples that contained significant cellular debris, making stage determination difficult. Cycle monitoring was performed in two cohorts of female WT and *Scn2a*^*E/+*^ mice beginning at 5-8 weeks of age. The first cohort consisted of mice that underwent either OVX or sham surgery (*n* = 19–20 per genotype and treatment, blinded to treatment). Subjects were allowed 7-14 days to recover from surgery before daily estrous cycle monitoring was performed leading up to, and on the day of, flurothyl seizure induction (**Figure 1A**). The monitoring period was at least 9 days for each for each subject, corresponding to approximately 2 cycles of average length^32^. Estrous cycle monitoring data was used to make blinded calls of surgical condition based on estrous cyclicity (i.e., cycle regularity). A subject was defined as having an irregular cycle if cycle length (average time to progress from one stage of estrus to the next) exceeded 7 days or if the subject spent more than 50% of time in any given stage^24,33^. A subject was blind called as having undergone OVX if it had an irregular cycle with a majority of samples determined to be “unclear”^34^. A single subject was excluded from analyses due to being blind called as receiving a sham surgery despite actually receiving OVX. The second cohort consisted of consisted of non-surgerized mice (*n* = 20–21 per genotype) evaluated separately from the previous cohort. Daily estrous cycle monitoring was performed leading up to, and on the day of flurothyl seizure induction. The monitoring period was 12 days for each for each subject, corresponding to approximately 3 cycles of average length^32^. Estrous cycle monitoring data was used to evaluate estrous cyclicity using the criteria defined above. Detailed estrous cycle monitoring data is available in Supplementary Table S1.

**Figure 1.**
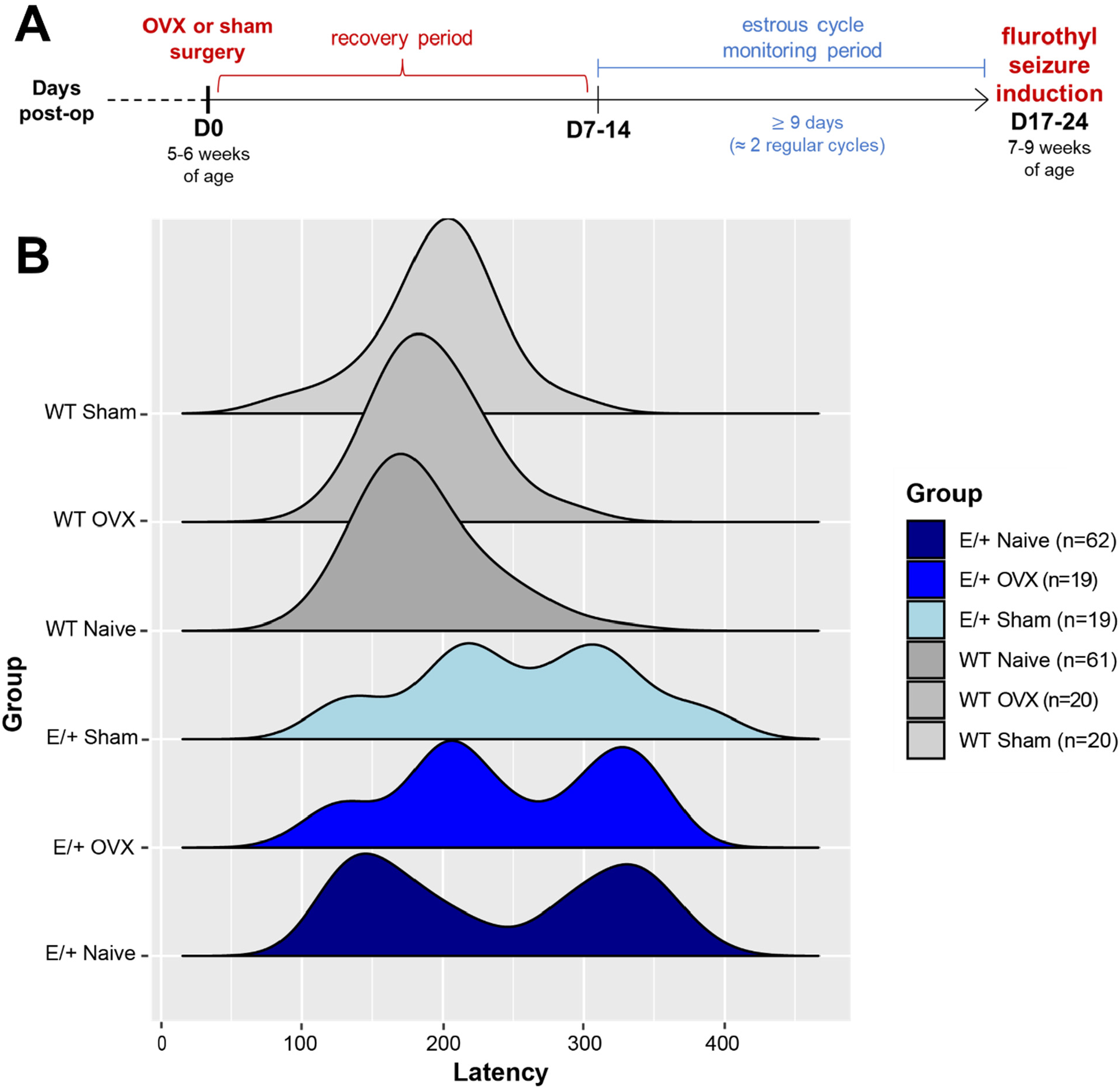
Ovariectomy does not abolish non-unimodal distribution of flurothyl-induced GTCS in *Scn2a*^*E/+*^ female mice. (**A**) Schematic of experimental design for time points of surgery, estrous cycle monitoring and flurothyl seizure induction. Ovariectomy is abbreviated as OVX. (**B**) Latency to first GTCS in WT and *Scn2a*^*E/+*^ female mice under different surgical conditions. Previously published data^6^ are replotted as WT Naïve and E/+ Naïve. Distributions of latencies to GTCS were evaluated for unimodality/non-unimodality using hypothesis testing (**Table 1**). Data from naïve *Scn2a*^*E/+*^ females had a non-unimodal distribution while naïve WT females had a unimodal distribution. WT females that underwent either OVX or sham surgery also had unimodal distributions. *Scn2a*^*E/+*^ females that underwent OVX had a non-unimodal distribution similar to naïve controls. *Scn2a*^*E/+*^ females that underwent sham surgery had a distribution that did not reach statistical significance for non-unimodality but is still significantly different from WT controls (*P=*0.0218, Kolmogorov-Smirnov test).

### Flurothyl seizure induction

Susceptibility to seizures induced by the chemoconvulsant flurothyl (Bis(2,2,2-trifluoroethyl) ether, Sigma-Aldrich, St. Louis, MO, USA) was assessed in two cohorts of female WT and *Scn2a*^*E/+*^ mice at 7-10 weeks of age. Flurothyl was introduced into a clear, plexiglass chamber (2.2 L) by a syringe pump at a rate of 20 µL/min and allowed to volatilize. Latency to first GTCS with loss of posture was recorded. The first cohort consisted of mice that underwent either OVX or sham surgery (*n* = 19–20 per genotype and treatment). The second cohort consisted of non-surgerized mice (*n* = 20–21 per genotype) evaluated separately from the previous cohort.

**Table 1.** Statistical comparisons

| Figure | Comparison | Test | Value | Post Hoc |
| --- | --- | --- | --- | --- |
| 3.3 | WT Sham GTCS Distribution | Hartigan’s Dip Test | p=0.4102 | n/a |
|  |  | Excess Mass Test | p=0.208 | n/a |
|  | WT OVX GTCS Distribution | Hartigan’s Dip Test | p=0.8703 | n/a |
|  |  | Excess Mass Test | p=0.624 | n/a |
|  | WT Naïve GTCS Distribution | Hartigan’s Dip Test | p=0.9935 | n/a |
|  |  | Excess Mass Test | p=0.986 | n/a |
|  | *Scn2a*<sup>E/+</sup> Sham GTCS Distribution | Hartigan’s Dip Test | p=0.4102 | n/a |
|  |  | Excess Mass Test | p=0.188 | n/a |
|  | *Scn2a*<sup>E/+</sup> OVX GTCS Distribution | Hartigan’s Dip Test | p=0.0479 | n/a |
|  |  | Excess Mass Test | p=0.02 | n/a |
|  | *Scn2a*<sup>E/+</sup> Naïve GTCS Distribution | Hartigan’s Dip Test | p=0.0006 | n/a |
|  |  | Excess Mass Test | p<0.0001 | n/a |
| 3.4 | Sham – Estrous Cycle Stage vs GTCS (A) | Two-way ANOVA | F(1,31)=1.552, p=0.2221<br>(Main Effect: Estrous Cycle Stage)<br>F(1,31)=9.791, p=0.0038<br>(Main Effect: Genotype) | Sidak’s |
|  | Sham Estrous Cycle Regularity (B) | Fisher’s Exact test | p=0.2003 | n/a |
| 3.5 | Untreated – Estrous Cycle Stage vs GTCS (A) | Two-way ANOVA | F(1,32)=0.7885, p=0.3812<br>(Main Effect: Estrous Cycle Stage)<br>F(1,32)=18.47, p=0.0002<br>(Main Effect: Genotype) | Sidak’s |
|  | Untreated Estrous Cycle Regularity (B) | Fisher’s Exact test | p=0.0109 | n/a |

### Statistical analysis

**Table 1** summarizes statistical tests used for all comparisons along with computed values. Distributions of latencies to flurothyl-induced GTCS were evaluated for unimodality/non-unimodality in six groups based on genotype and surgical condition using the R packages ‘diptest’ and ‘multimode’ (RStudio 4.2.0)^35–37^. Two distributions representing previously published data^6^ from untreated female WT and *Scn2a*^*E/+*^ mice are replotted as “WT Naïve” (*n* = 61) and “E/+ Naïve” (*n* = 62) in **Figure 1B**. These distributions were not previously evaluated specifically for unimodality/non-unimodality. Mice from these historical groups did not undergo estrous cycle monitoring. The remaining four distributions represent WT and *Scn2a*^*E/+*^ mice that underwent either OVX or sham surgery and subsequent estrous cycle monitoring. All other comparisons were performed as indicated in **Table 1** using GraphPad Prism (v 9.4.1).

## Results & Discussion

### Ovariectomy does not abolish non-unimodal distribution of flurothyl seizure thresholds in Scn2a^E/+^ mice

To evaluate whether the estrous cycle is associated with flurothyl seizure thresholds in *Scn2a*^*E/+*^ mice, we performed ovariectomy (OVX) or sham surgery. A schematic of the experimental design is shown in **Figure 1A**. After a recovery period of 7-14 days, we performed daily monitoring of the estrous cycle. Flurothyl was used to induce GTCS in WT and *Scn2a*^*E/+*^ mice 17-24 days after surgery (7-9 weeks of age). Specifically, we wanted to compare the distribution of latencies to GTCS in surgically treated mice against historical data from untreated mice^6^. We reasoned that if circulating sex hormones were affecting flurothyl-seizure threshold, then removing said hormones via OVX would collapse the distribution observed in *Scn2a*^*E/+*^ females such that it would be more similar to the distributions observed in WT females or *Scn2a*^*E/+*^ males^6^. In order to statistically evaluate distributions for unimodality/non-unimodality, we used Hartigan’s *dip* test and the multimode test proposed by Ameijeiras-Alonso and colleagues^38,39^.

Both analytical methods involve hypothesis testing in which unimodality represents the null hypothesis and a *P*-value of ≤ 0.05 resulted in rejection of the null hypothesis. *P*-values for all groups are shown in **Table 1**. The historical data from untreated (naïve) *Scn2a*^*E/+*^ females had a non-unimodal distribution, while historical data from naïve WT females had a unimodal distribution (**Figure 1B**). Surgically treated (OVX and sham) WT females also had unimodal distributions of GTCS latencies similar to naïve controls (**Figure 1B**). Contrary to our hypothesis, OVX did not affect the distribution of GTCS latencies in *Scn2a*^*E/+*^ females; the distribution representing this group was non-unimodal (**Figure 1B**). The above data suggest that estrous cyclicity does not significantly contribute to flurothyl seizure thresholds in WT or *Scn2a*^*E/+*^ mice. *Scn2a*^*E/+*^ females that underwent sham surgery had a distribution that did not reach statistical significance for non-unimodality, although it was still significantly different from WT controls as determined by Kolmogorov-Smirnov test comparing cumulative distributions (**Figure 1B**, *P=*0.0218).

### Flurothyl seizure threshold is not associated with estrous cycle stage in Scn2a^E/+^ mice

Estrous cycle monitoring in surgically treated mice served a dual purpose. First, it allowed us to evaluate the success of OVX versus sham surgeries. Blind calls of surgical condition based on estrous cycle monitoring were nearly 100% accurate, with the exception of a single subject which was excluded. Second, it allowed us to evaluate whether flurothyl seizure threshold is associated with a particular stage of the estrous cycle in *Scn2a*^*E/+*^ mice that underwent sham surgery. We grouped subjects based on the estrous cycle stage determined on the day of flurothyl seizure induction. For the purposes of analysis, we compared stages that are estradiol-dominant (proestrus and estrus) with diestrus (progesterone-dominant)^24,40^. Two-way ANOVA comparing average latency to GTCS in sham surgery *Scn2a*^*E/+*^ females and WT controls showed only a significant main effect of genotype, but no significant effect of estrou cycle stage or genotype-by-stage interaction (**Table 1**) Although the genotype effect appears to be driven by the pro/estrus groups such that *Scn2a*^*E/+*^ females had a higher threshold for GTCS (271.0 ± 67.0 s) compared to WT (201.3 ± 45.2 s; **Figure 2A**), the WT diestrus group was underrepresented (n = 4). The observed genotype effect is also in agreement with our previous findings on flurothyl seizure threshold in *Scn2a*^*E/+*^ mice^6^ This data, along with lack of observable effects in the OVX condition, provides convergent evidence that the estrous cycle does not significantly affect flurothyl seizure thresholds in *Scn2a*^*E/+*^ mice Other neurobiological mechanisms apart from the acute influence of sex hormones have been proposed to address sex differences in seizure susceptibility^10^. In the case of flurothyl seizure threshold in *Scn2a*^*E/+*^ mice, any proposed mechanism would have to account for why the distribution of seizure thresholds in uniquely non-unimodal in *Scn2a*^*E/+*^ females. One possibility is that variation in one or more modifier genes confers sex-specific protection to females such that there is an increased threshold for phenotypic penetrance of the K1422E variant. This is similar to a hypothesis that has been proposed to explain the “female protective effect” in ASD^9,41–43^. Another possibility is that female mice are differentially exposed to androgens *in utero* based on their intrauterine position relative to male siblings^44^. Androgen exposure *in utero* is known to mediate early brain development and could influence seizure susceptibility in differentially exposed females^10,45^.

**Figure 2.**
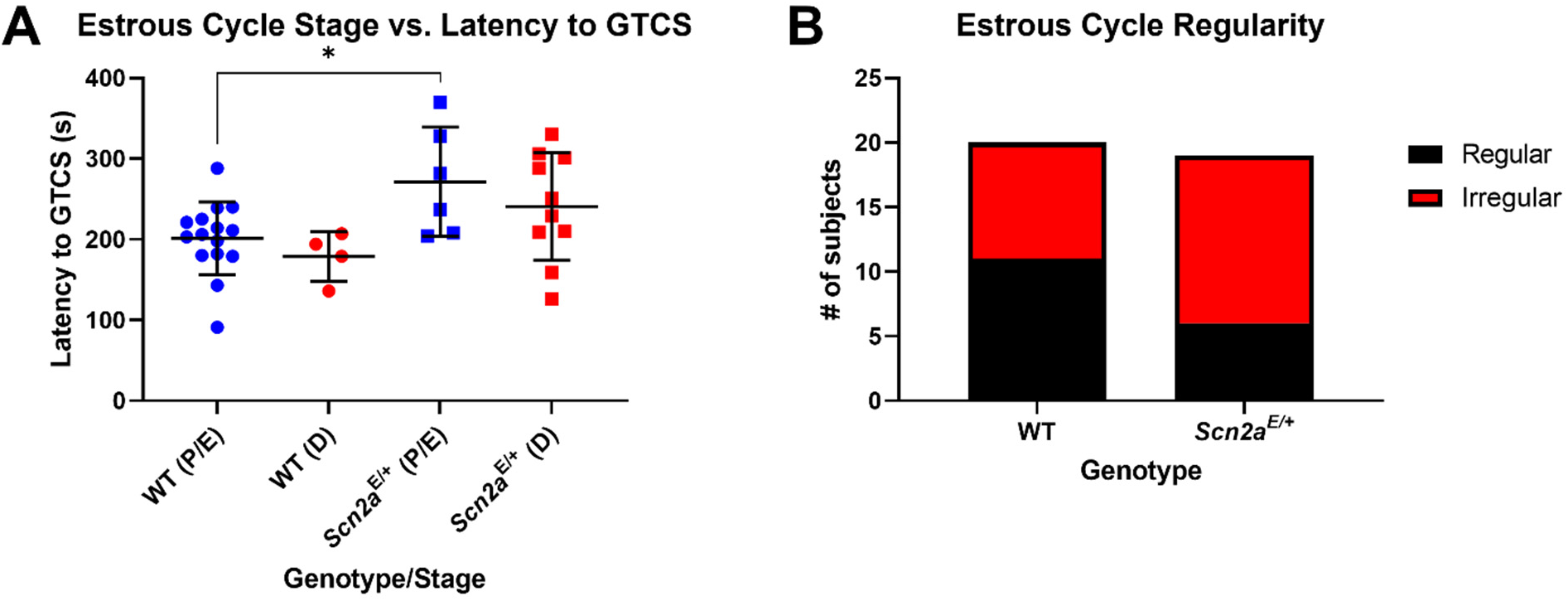
Latency to flurothyl-induced GTCS is not associated with estrous cycle stage in sham ovariectomized *Scn2a*^*E/+*^ female mice. (**A**) Latency to first flurothyl-induced GTCS in sham ovariectomized WT and *Scn2a*^*E/+*^ female mice during pro/estrus (abbreviated P/E) or diestrus (abbreviated D). Two-way ANOVA showed only a significant main effect of genotype [*F*(1,31)=9.791, *P*=0.0038]. Post hoc-analysis showed that *Scn2a*^*E/+*^ females in pro/estrus had an elevated threshold for GTCS compared to stage-matched WT controls (WT: 201.3 ± 45.2 s, *Scn2a*^*E/+*^: 271.0 ± 67.0 s, ^*^*P=*0.0268; Sidak’s post-hoc test). Symbols represent individual mice, horizontal lines represent mean, and error bars represent SD; *n* = 4-15/stage/genotype. (**B**) Number of sham ovariectomized WT and *Scn2a*^*E/+*^ female mice (*n =* 19-20 per genotype) with regular vs irregular estrous cycles (WT: 45.0% irregular, *Scn2a*^*E/+*^:68.4% irregular). Proportion of mice with irregular cyclicity was not significantly different between genotypes (*P=*0.2003, Fisher’s Exact test). Irregular cyclicity was defined as having a cycle length (average time to progress from one stage of estrus to the next) that exceeded 7 days as spending more than 50% of the monitoring period in any given stage.

### Disrupted estrous cyclicity in Scn2a^E/+^ mice

As part of our estrous cycle monitoring in surgically treated mice, we evaluated overall cycle regularity. A subject was defined as having an irregular cycle if cycle length exceeded 7 days or if the subject spent more than 50% of time in any given stage^24,33^. We noted that a proportion of both WT and *Scn2a*^*E/+*^ sham ovariectomized mice had irregular cyclicity (WT: 45.0%, *Scn2a*^*E/+*^: 68.4%), likely because of surgical intervention (**Figure 2B**). In order to exclude the confounding factor of surgical trauma, we wanted to evaluate whether flurothyl seizure threshold is associated with a particular stage of the estrous cycle in an additional cohort of untreated mice. Similar to what was observed in sham surgery mice (**Figure 2A**), two-way ANOVA comparing average latency to GTCS in untreated *Scn2a*^*E/+*^ females and WT controls showed only a significant main effect of genotype but no significant effect of estrus cycle stage or genotype-by-stage interaction (**Table 1**). Again, this effect appears to be driven by the pro/estrus groups such that *Scn2a*^*E/+*^ females had a higher threshold for GTCS (290.0 ± 67.0 s) compared to WT (185.5 ± 30.6 s; **Figure 3A**). Interestingly, while the proportion of sham surgery mice with irregular cyclicity was not significantly different between genotypes (**Figure 2B**), the proportion of untreated *Scn2a*^*E/+*^ females with irregular cyclicity was significantly greater than WT (WT: 19.1%, *Scn2a*^*E/+*^: 60.0% irregular; *P=*0.0109, Fisher’s exact test; **Figure 3B**). This suggests that disrupted estrous cyclicity may be causally associated with the K1422E variant.

**Figure 3.**
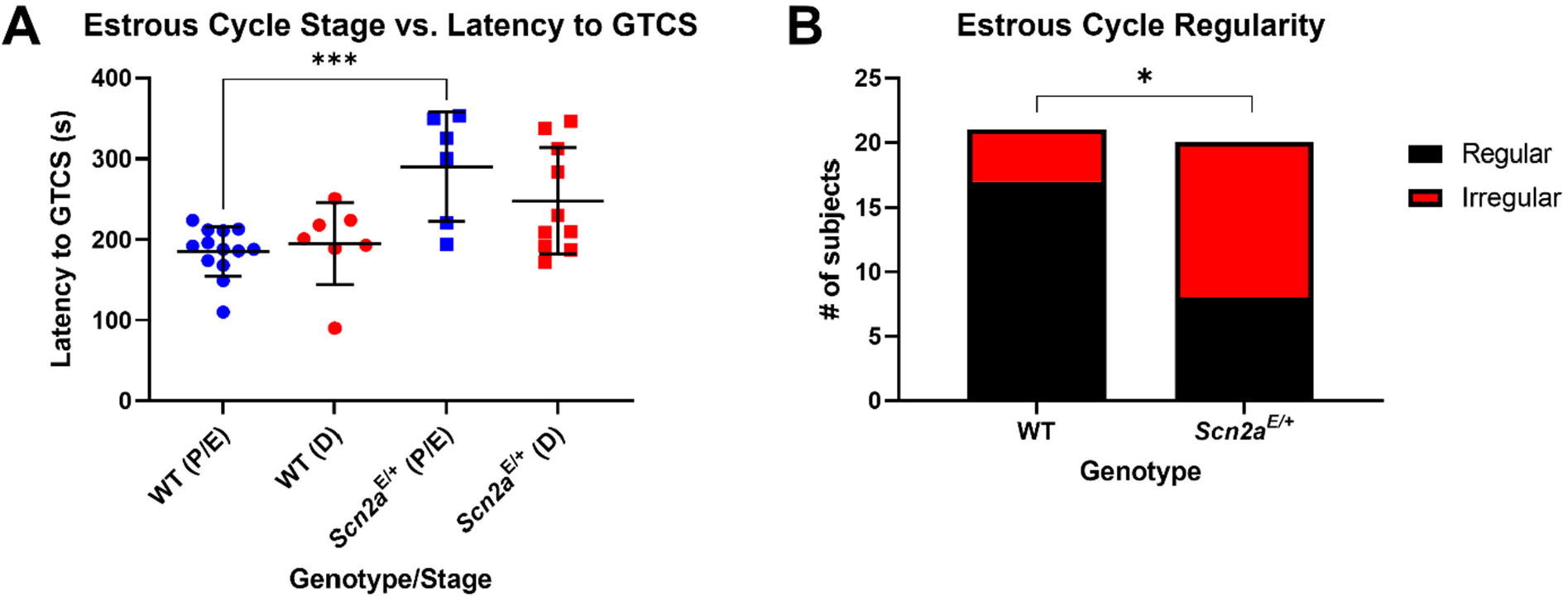
Disrupted estrous cyclicity in untreated *Scn2a*^*E/+*^ female mice. (**A**) Latency to first flurothyl-induced GTCS in untreated WT and *Scn2a*^*E/+*^ female mice during pro/estrus (abbreviated P/E) or diestrus (abbreviated D). Two-way ANOVA showed only a significant main effect of genotype [*F*(1,32)=18.47, *P*=0.0002]. Post hoc-analysis showed that *Scn2a*^*E/+*^ females in pro/estrus had an elevated threshold for GTCS compared to stage-matched WT controls (WT: 185.5 ± 30.6 s, *Scn2a*^*E/+*^: 290.0 ± 67.0 s, ^***^*P=*0.0006; Sidak’s post-hoc test). Symbols represent individual mice, horizontal lines represent mean, and error bars represent SD; *n* = 6-13/stage/genotype. (**B**) Number of sham untreated WT and *Scn2a*^*E/+*^ female mice (*n =* 20-21 per genotype) with regular vs irregular estrous cycles (WT: 19.1% irregular, *Scn2a*^*E/+*^:60.0% irregular). Proportion of *Scn2a*^*E/+*^ female mice with irregular cyclicity was significantly greater compared to WT (^*^*P=*0.0109, Fisher’s Exact test). Irregular cyclicity was defined as having a cycle length (average time to progress from one stage of estrus to the next) that exceeded 7 days as spending more than 50% of the monitoring period in any given stage.

Disrupted estrous cyclicity has been observed in other rodent models of epilepsy, but this is the first demonstration of such an effect in a pathogenic variant model, particularly *SCN2A*-related NDD^33,46–48^. This is important because epilepsy is generally associated with an increased risk for reproductive endocrine disorders, such as disrupted menstration^10,17,18^. However, the degree of comorbidity between these conditions and *SCN2A*^*-*^related NDD specifically has not been formally studied. It has been shown that circadian signals from the suprachiasmatic nucleus (SCN) to gonadotropin-releasing hormone (GnRH) neurons in the hypothalamus are required for estrous cyclicity^49,50^. Sleep disturbances are frequently reported in individuals with *SCN2A-* related NDD, highlighting a connection between *SCN2A* and circadian rhythms^51^. Region-specific deficiency of *Scn2a* has been used model sleep disturbances in mice and is associated with disrupted firing of SCN neurons and changes in the expression of circadian entrainment pathway genes^52^. This suggests a possible mechanism by which changes in *Scn2a* function could lead to disrupted estrous cyclicity.

## Limitations

A limitation of the current study is that *Scn2a*^*E/+*^ females in the sham surgery condition had a distribution that did not reach statistical significance for non-unimodality (**Figure 1B**). This observation, along with the non-zero proportion of both WT and *Scn2a*^*E/+*^ sham surgery mice that had irregular estrous cycles (**Figure 2B**), suggests a potentially confounding effect of surgical intervention. To address this potential limitation, we repeated estrous cycle monitoring and flurothyl seizure induction in an additional cohort of untreated mice. Another limitation is that group numbers were unequal when comparing estrous cycle stage because we opted to maximize the number of subjects undergoing seizure induction on the same day under identical conditions. However, due to the lack of effect in the OVX treatment condition, there was little incentive to round out these estrous stage group numbers (**Figure 2A & 3A**).

## Conclusions

Overall, we believe the current study reflects a technically sound experimental approach to probe the questions raised by our hypothesis, and our results support rejection of our original hypothesis that estrous cycle affects seizure susceptibility in *Scn2a*^*E/+*^ mice. Importantly, in conducting this study, we discovered disrupted estrus cyclicity in *Scn2a*^*E/+*^ mice, highlighting the value of investigating sex-specific effects and estrous cycle in genetic epilepsies and NDD, as well as in the broader field of neuroscience.

## Abbreviations

ASD: autism spectrum disorder
flurothyl: Bis(2,2,2-trifluoroethyl) ether
GTCS: generalized tonic-clonic seizure
GnRH: gonadotropin-releasing hormone
IP: intraperitoneal
NDD: neurodevelopmental disorders
OVX: ovariectomy
PBS: phosphate buffered saline
SC: subcutaneous
SCN: suprachiasmatic nucleus
SD: standard deviation

## CRediT authorship contribution statement

**Dennis M. Echevarria-Cooper:** Conceptualization, Methodology, Formal analysis, Investigation, Writing -Original Draft, Writing - Review & Editing, Visualization. **Jennifer A. Kearney:** Conceptualization, Formal analysis, Writing - Review & Editing, Project administration, Funding acquisition.

## Data Availability

Individual subject level data are available in Supplementary Table 1.

## Acknowledgements

We thank Dr. Emma Liechty for ovariectomy training and Nathan Speakes for technical assistance. This work was supported by the National Institutes of Health grant U54 NS108874 (JAK).

